# Oxidative stress resistance and fitness-compensatory response in vancomycin-intermediate *Staphylococcus aureus* (VISA)

**DOI:** 10.1101/297036

**Authors:** Xin-Ee Tan, Hui-min Neoh, Longzhu Cui, Keiichi Hiramatsu, Rahman Jamal

## Abstract

In this study, VISA cells carrying *vraS* and/or *graR* mutations were shown to be more resistant to oxidative stress. *Caenorhabditis elegans* infected with these strains in turn demonstrated lower survival. Altered regulation in oxidative stress response and virulence appears to be physiological adaptations associated with VISA phenotype in the Mu50 lineage.

---

Bacterial antibiotic resistance has been reported to occur concurrently with changes in various cellular responses of the organism. In particular, altered virulence mechanism is common among antibiotic resistant strains. Acquisition of antibiotic resistance often imposes a fitness burden on bacterial cells (1); in most cases, increased resistance has been paralleled with decreased virulence, as reported in methicillin-resistant *Staphylococcus aureus* (2, 3) and vancomycin-intermediate *S. aureus* (VISA) (4, 5). Apart from virulence, the association between antibiotic resistance and oxidative stress response has also been reported. Different classes of antibiotics, regardless of their primary targets, have been shown to induce lethality through generation of reactive oxygen species (ROS) (6, 7). In response, the bacteria will try to reduce antibiotic killing via reduction of cellular hydroxyl radical accumulation (8–12).

We previously employed a proteomic approach to determine underlying regulatory pathway(s) mediating transition of vancomycin-susceptible *S. aureus* (VSSA, strain Mu50Ω) to VISA (strain Mu50Ω-*vraS*m, harbouring a *vraS* T700A mutation; and strain Mu50Ω-*vraS*m-*graR*m, harbouring both *vraS* T700A*/graR* A590G mutations compared to strain Mu50Ω) (13). In the study, unexpected features of up-regulated oxidized protein repair enzyme (MsrB) and down-regulated virulence-associated proteins (Spa, Rot, MgrA, SarA) in VISAs were observed. Functional categorization and differential proteomic profiles of total proteins extracted from the 3 isogenic strains are presented in Figure 1 and Figure 2, respectively. Consistent up-regulation of MsrB as well as down-regulation of virulence-associated proteins in VISA strains lead us to suspect possible interplay between oxidative stress response, virulence and antibiotic resistance in VISA strains of the Mu50 lineage.

**Figure 1:**
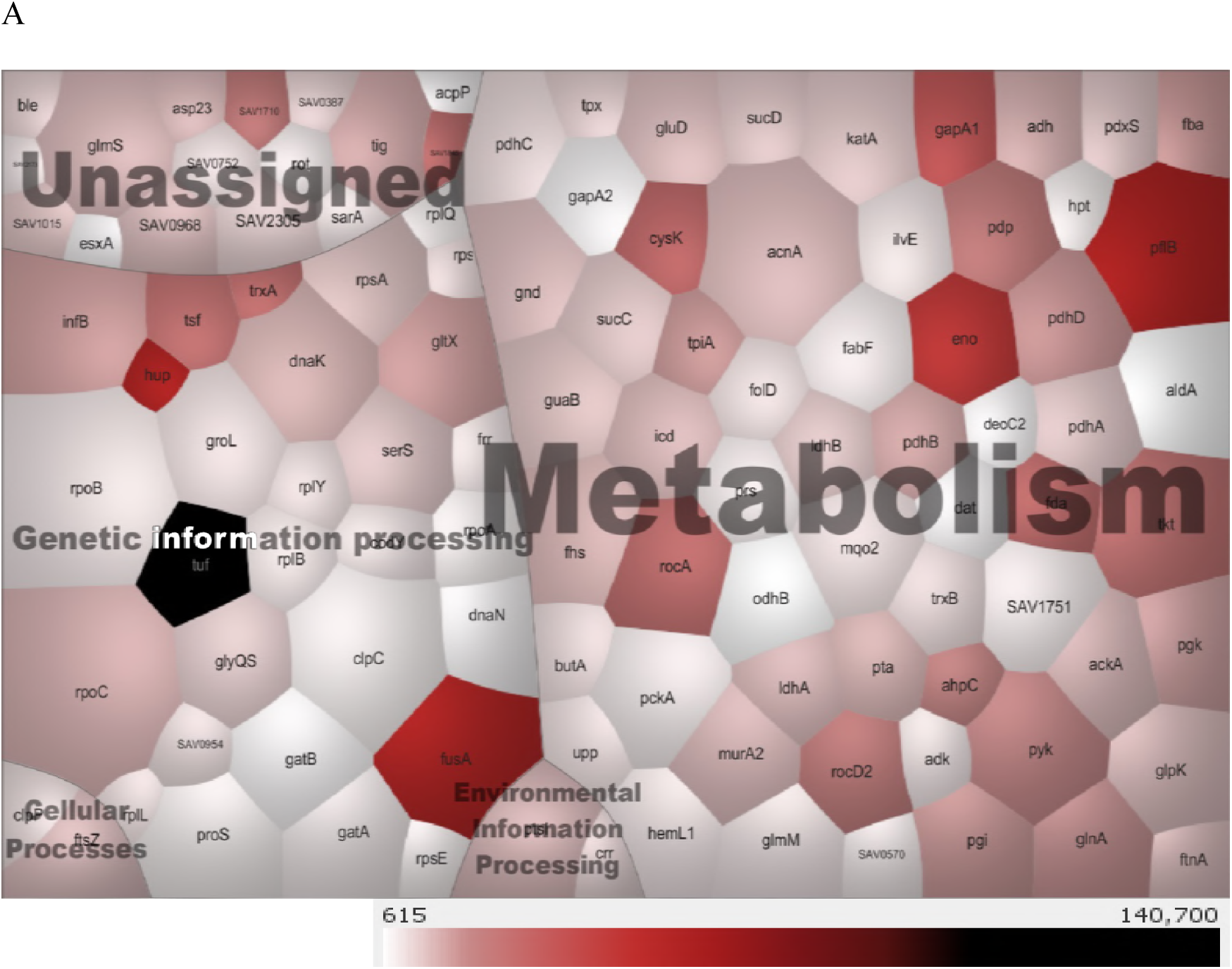
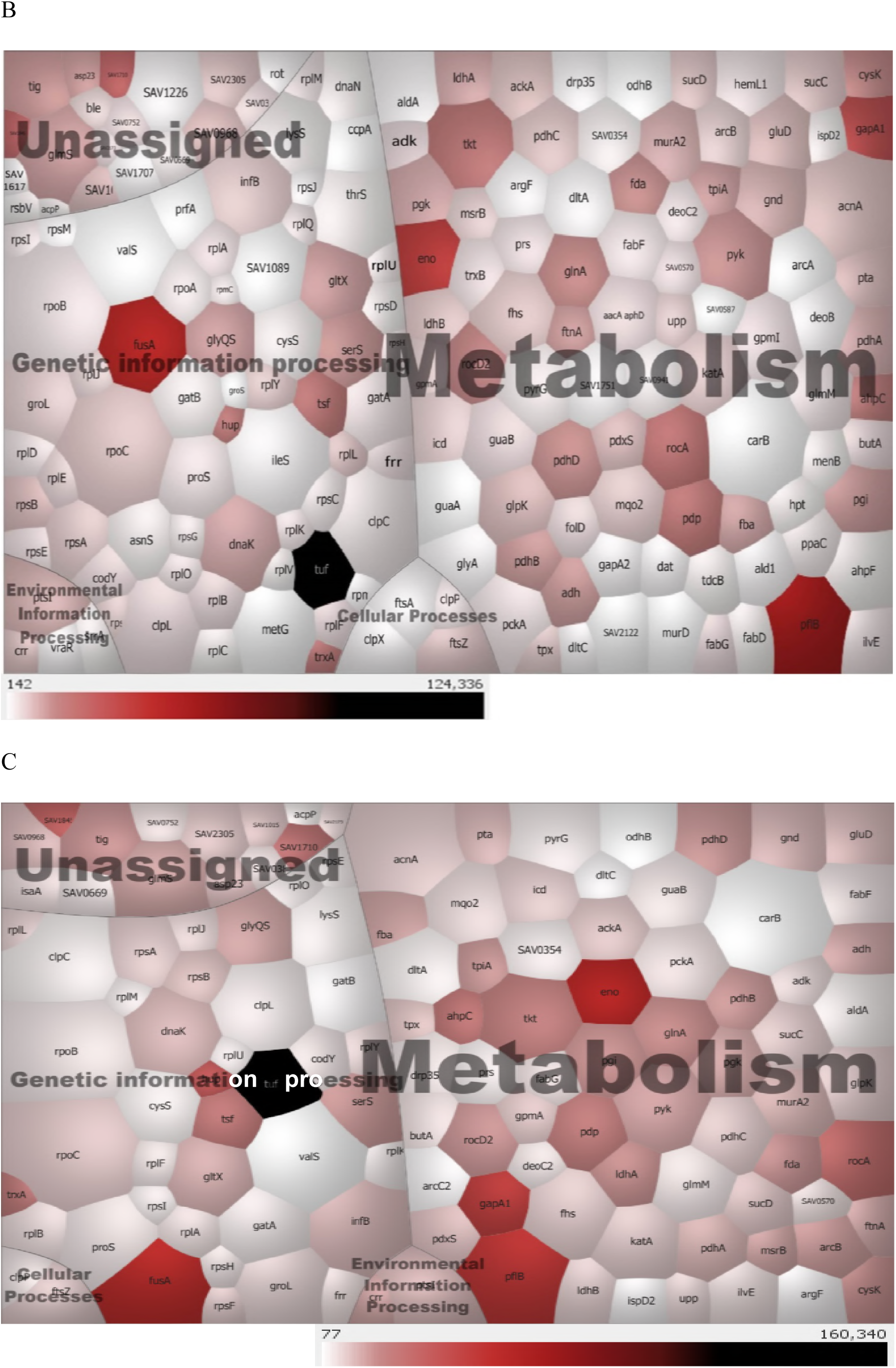
Voronoi mapping of total proteins extracted from Mu50Ω (panel A), Mu50Ω-*vraS*m (panel B) and Mu50Ω-*vraS*m-*graR*m (panel C). Each cell in the voronoi treemap represents one protein. Colour intensity of each cell is proportional to its protein abundance while cell size is relative to protein chain length. Total proteins have been categorized into 5 groups, with the majority of proteins found to be involved in cellular metabolism and genetic information processing.

**Figure 2:**
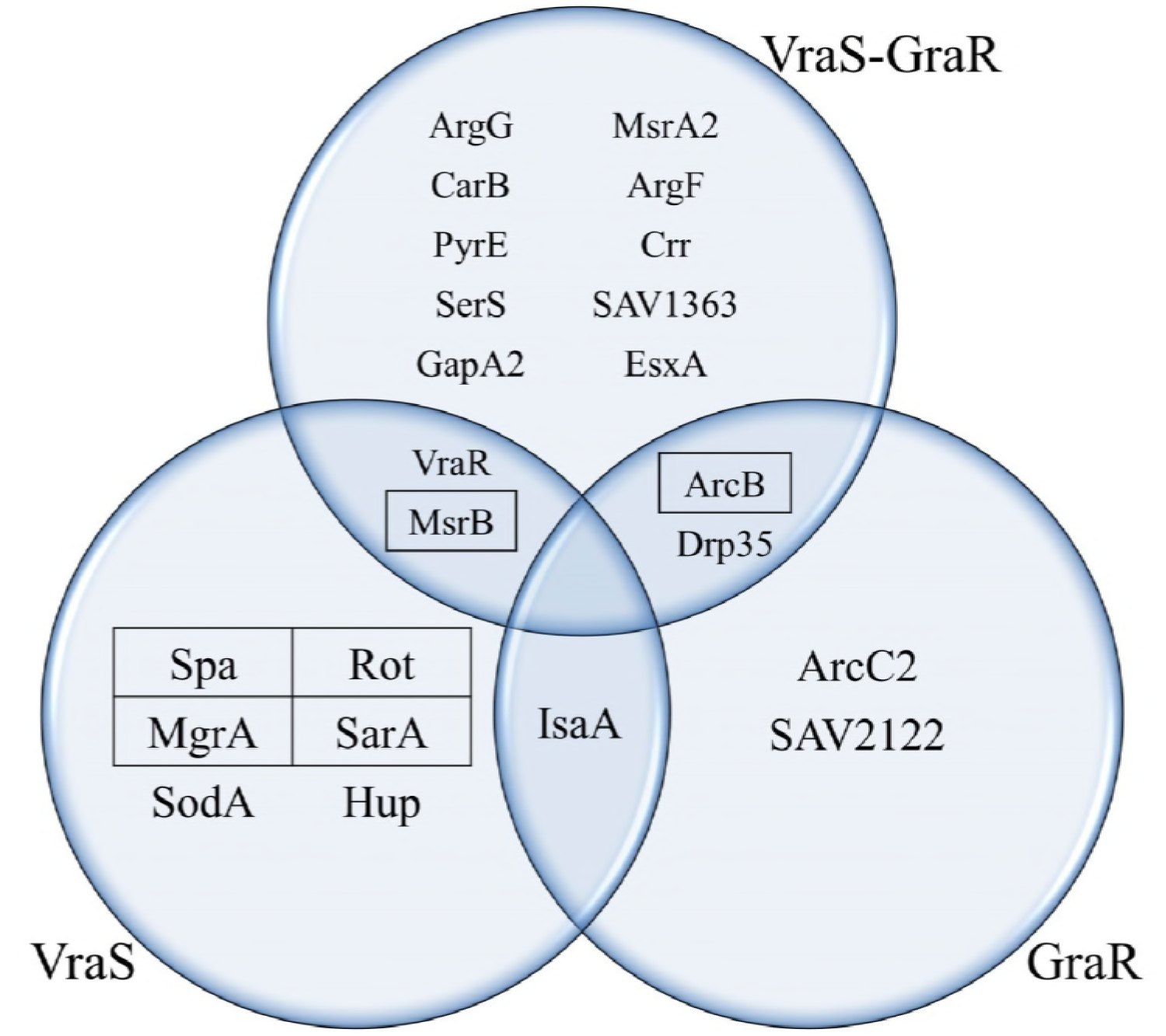
Comparative proteomic profiling of Mu50Ω, Mu50Ω-*vraS*m and Mu50Ω-*vraS*m-*graR*m revealed differential protein expression profiles regulated by the VraS, GraR and VraS-GraR regulons (comparison of protein profiles between Mu50Ω-*vraS*m and Mu50Ω, Mu50Ω-*vraS*m-*graR*m and Mu50Ω-*vraS*m, and between Mu50Ω-*vraS*m-*graR*m and Mu50Ω, respectively) (13). Virulence-related proteins (Spa, Rot, MgrA, SarA), MsrB and ArcB (proteins in text boxes) are the proteins of interest selected for further investigation of their association with vancomycin resistance as they were found to be differentially expressed in VISAs compared to VSSA.

Methionine sulfoxide reductases (Msr) are bacterial repair enzymes important for protection from oxidative killing (14–17). To determine if different MsrB expression levels in the 3 study strains affect their responses towards oxidative stress, Mu50Ω, Mu50Ω-*vraS*m and Mu50Ω-*vraS*m-*graR*m were treated with 3 oxidizing agents [cumene hydroperoxide, tert-butyl hydroperoxide and hydrogen peroxide (H_2_O_2_)] at various concentrations prior determination of viable cell counts. Interestingly, Mu50Ω-*vraS*m and Mu50Ω-*vraS*m-*graR*m were shown to have greater survival when challenged with cumene hydroperoxide (Figure 3) and tert-butyl hydroperoxide (Figure 4) compared to Mu50Ω, indicating that VISA strains (with up-regulated MsrB) exhibited higher resistance towards oxidative damage.

**Figure 3:**
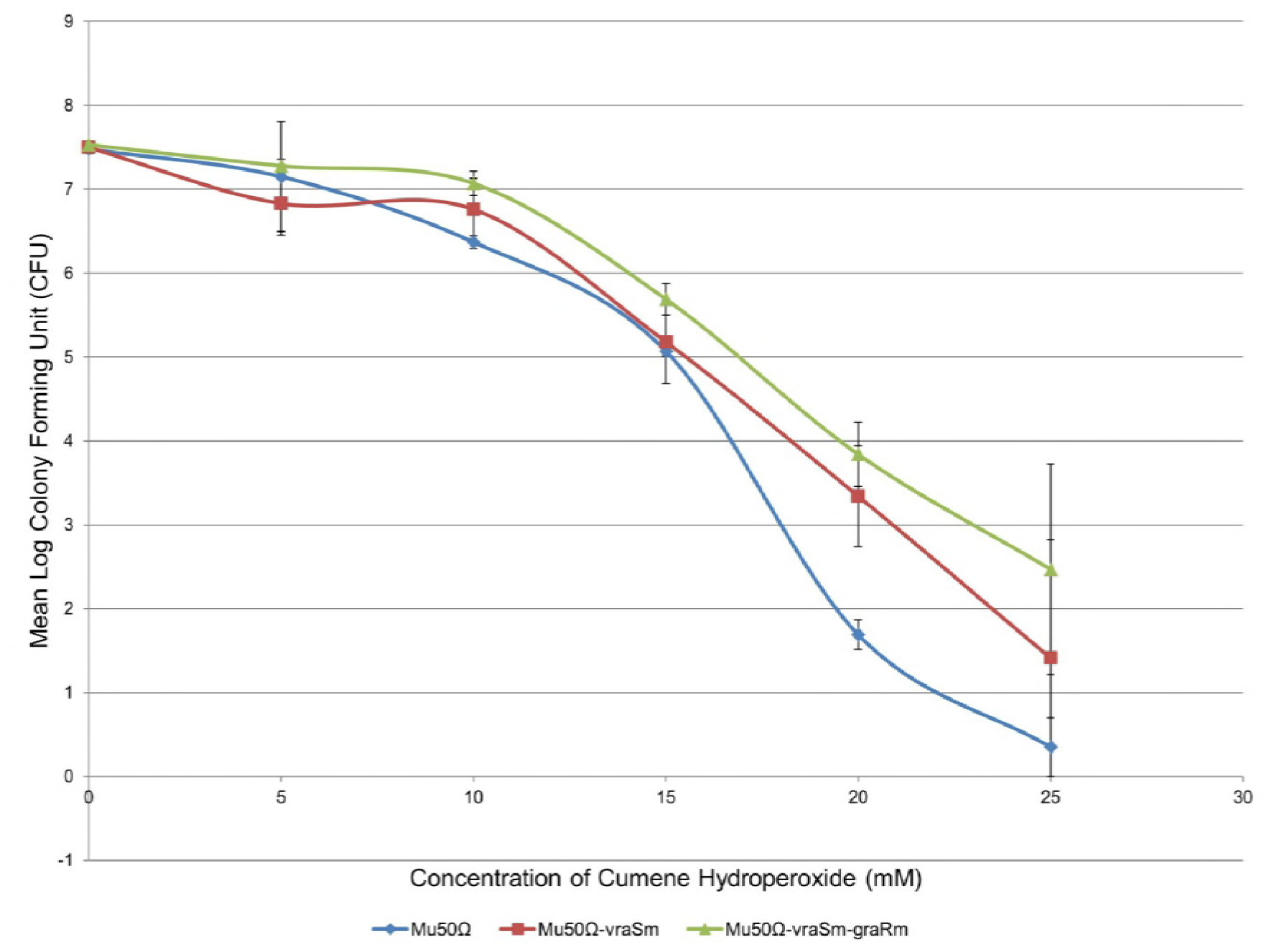
Cumene hydroperoxide oxidative stress test on Mu50Ω, Mu50Ω-*vraS*m and Mu50Ω-*vraS*m-*graR*m. Both VISA strains showed higher resistance to oxidative stress compared to VSSA.

**Figure 4:**
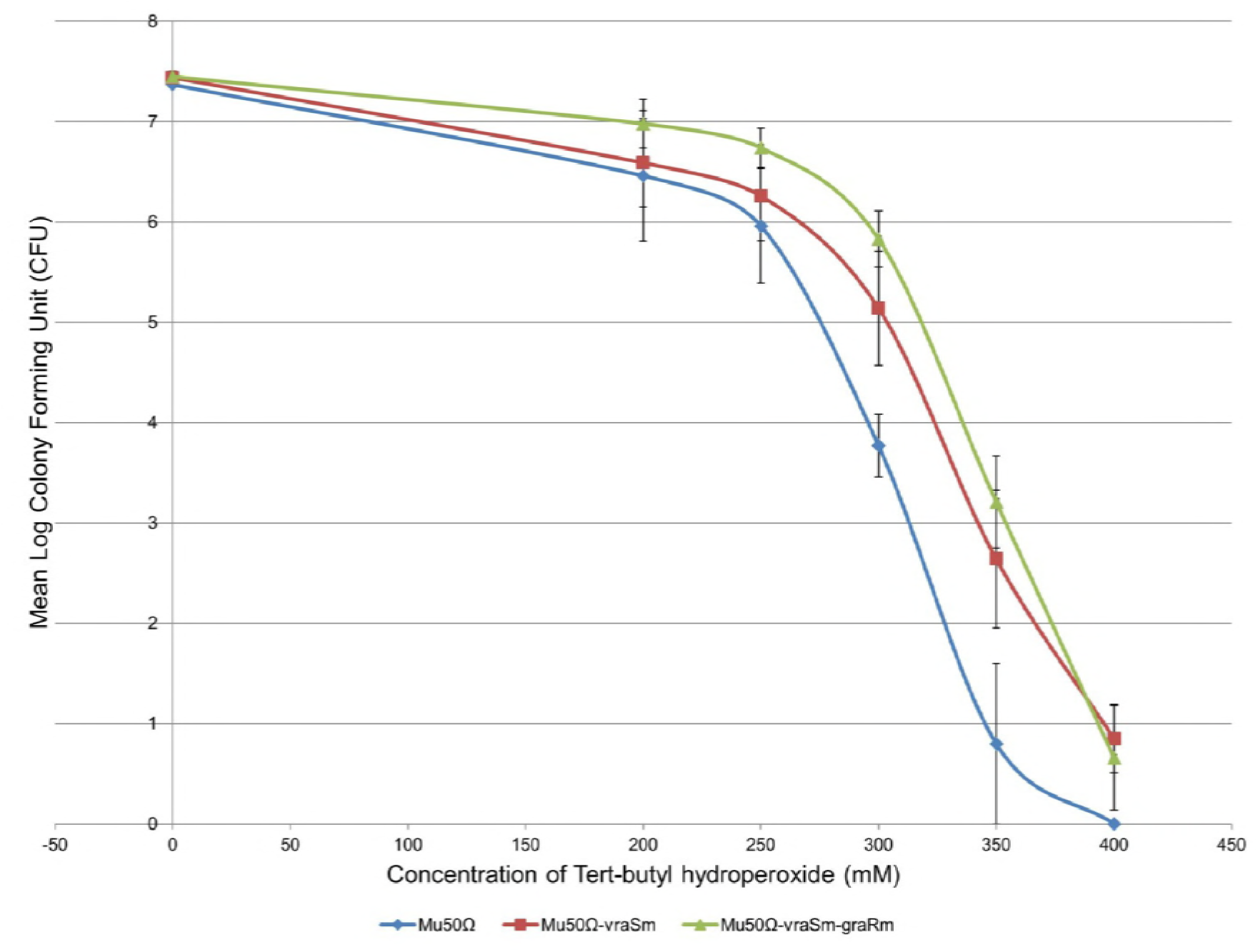
Tert-butyl hydroperoxide oxidative stress test on Mu50Ω, Mu50Ω-*vraS*m and Mu50Ω-*vraS*m-*graR*m. Both VISA strains were more resistant to oxidative killing compared to VSSA.

We postulate that in order to circumvent oxidative damage caused by vancomycin (6), VISA cells are primed to produce Msr proteins, which are the only known enzymes capable of reducing oxidized form of methionine, thereby restoring normal function of proteins (18). This cellular response is proposed to be mediated by VraSR system, since, in our previous study, up-regulation of MsrB proteins was identified in the VraS and VraS-GraR regulons, but not the GraR regulon (Figure 2). Accordingly, Pang et al.’s study demonstrated that complementation of *S. aureus vraSR* knockout mutant (Δ*vraSR*) restored its *msrA1* expression to a higher level compared with Δ*vraSR* (19).

Nevertheless, a different survival trend was observed in H_2_O_2_-induced VISA cells. Although having lower MsrB expression, VSSA Mu50Ω displayed greater survival after H_2_O_2_ induction compared with VISA strains (Figure 5). Increased susceptibility to H_2_O_2_ killing was previously reported to be associated with the lack of staphyloxanthin (carotenoid pigmentation) (20, 21). As suggested by Singh et al., *msrB* deletion reduced *S. aureus* susceptibility to H_2_O_2_, and this phenotype is accompanied by increased production of carotenoids in the mutant cells (22). In concordance, lower expression of MsrB in Mu50Ω might have resulted in higher levels of cellular carotenoids and subsequent resistance to H_2_O_2_.

**Figure 5:**
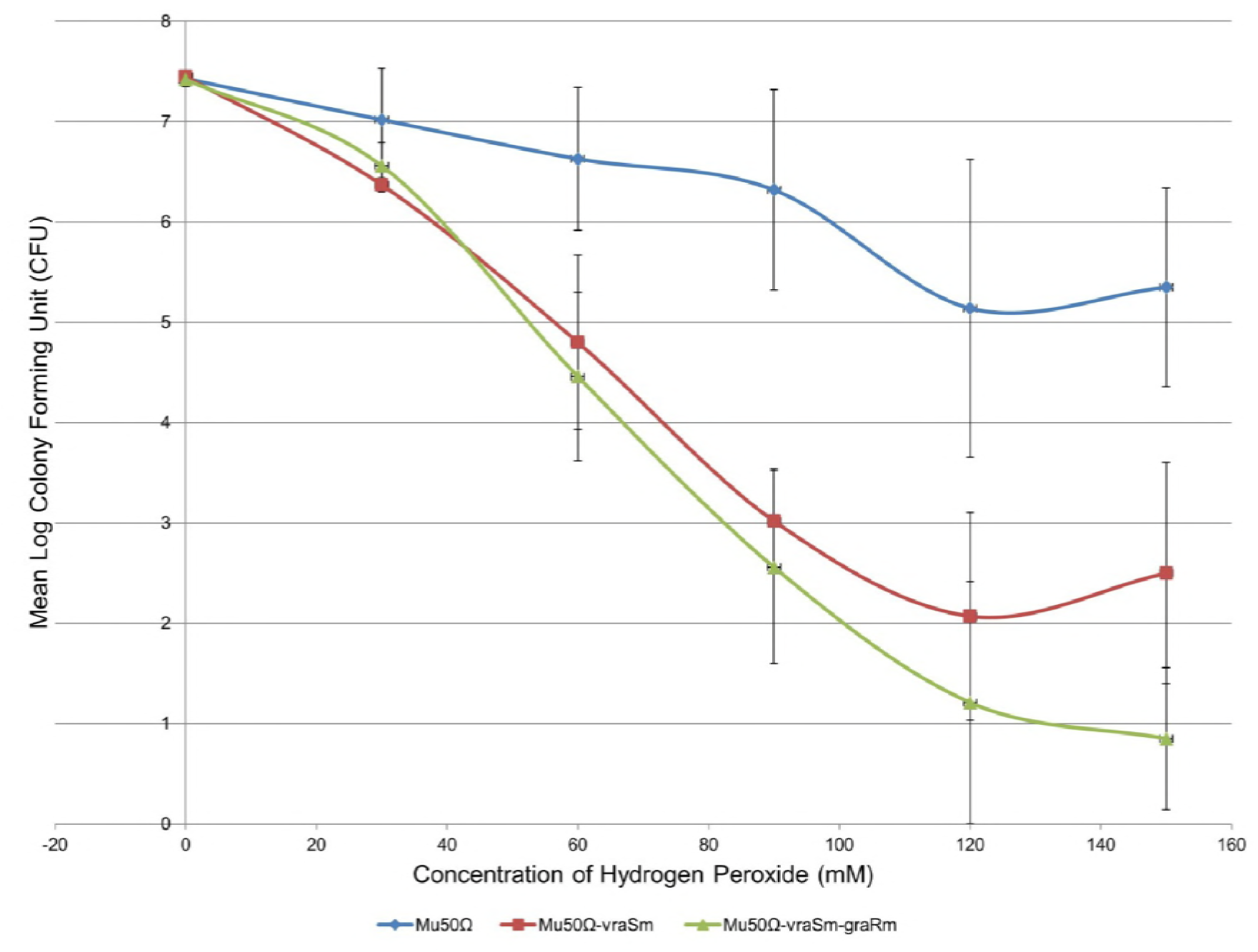
Hydrogen peroxide oxidative stress test on Mu50Ω, Mu50Ω-*vraS*m and Mu50Ω-*vraS*m-*graR*m. Mu50Ω was more resistant to oxidative stress from hydrogen peroxide induction compared to VISA strains.

In addition to increased resistance to oxidative stress, down-regulation of virulence-related proteins is also observed in the Mu50Ω-*vraS*m and Mu50Ω-*vraS*m-*graR*m VISAs (Figure 2) (13). We subsequently used a *Caenorhabditis elegans* survival assay to determine our study strains’ virulence (23). Forty L4 nematodes of *pos-1*-silenced *C. elegans* N2 strain were fed with the study strains of Mu50Ω, Mu50Ω-*vraS*m and Mu50Ω-*vraS*m-*graR*m, respectively; worm survival (quantity of live and dead worms) for every strain was then scored every 24 hours for 14 days and plotted on a Kaplan-Meier survival plot (Figure 6). The experiment showed that VISA strains exhibited higher nematocidal activity via complete killing of all 40 *C. elegans* on the 3^rd^ (Mu50Ω-*vraS*m-*graR*m) and 8^th^ (Mu50Ω-*vraS*m) day of the assay. On the other hand, killing of *C. elegans* fed with VSSA was gradual and surviving worms were still observed at the end of the 14-days assay.

**Figure 6:**
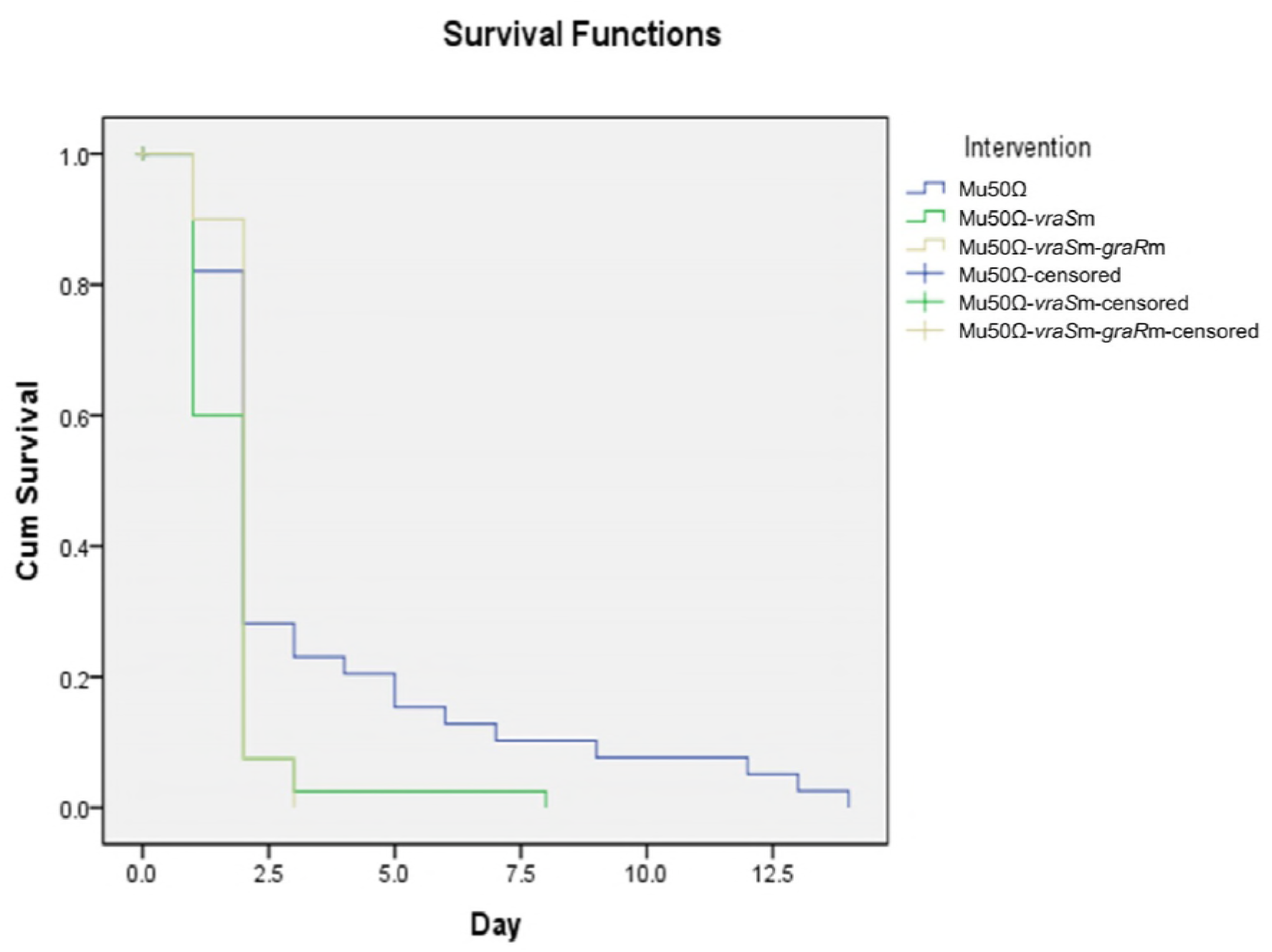
Kaplan-Meier survival plot for *C. elegans* fed with Mu50Ω, Mu50Ω-*vraS*m and Mu50Ω-*vraS*m-*graR*m. A significant decrease in survival of *C. elegans* infected with VISA strains Mu50Ω-*vraS*m (*p* < 0.05) and Mu50Ω-*vraS*m-*graR*m (*p* < 0.05) was observed compared to those infected with VSSA strain Mu50Ω. Pairwise comparison also demonstrated a significant reduction in the survival of Mu50Ω-*vraS*m-*graR*m-infected *C. elegans* compared with Mu50Ω-*vraS*m-infected worms (*p* < 0.05).

*C. elegans* exhibits specific immune response towards different infective microorganisms; transcription profiles of *C. elegans* exposed to *Candida albicans* has been shown to be different from those infected with *Pseudomonas aeruginosa* or *S. aureus* (24). Both living and heat-killed *S. aureus* have been reported to be capable of triggering *C. elegans* responses (25). These studies suggest that *C. elegans* distinguish infections from different pathogens via recognition of specific bacterial pathogen-associated molecular patterns (PAMPs). Spa, a *S. aureus* cell wall surface protein, has been reported to be one of the PAMPs found in this Gram-positive bacterium (26). In our study, we postulate that down-regulation of Spa protein in VISA strains diminished the capability of *C. elegans* innate immune system to identify the bacteria, allowing VISA to achieve immune evasion. Consequently, VISA infections of *C. elegans* were found to be more lethal compared with VSSA. Even though *C. elegans* produces ROS in response to *S. aureus* infection (27), as VISA strains in this study were found to be more resistant to oxidative killing due to higher expression of MsrB enzymes, the strains had a survival edge from the ROS attack of *C. elegans* compared to VSSA. This allows VISA strains to bypass *C. elegans* defence mechanisms, resulting in expedited killing of the hosts.

Taking into consideration the results from our previous (13, 28) and current studies, we propose the interplay between cellular metabolism, oxidative stress response and virulence in VISA strains of Mu50 lineage (Figure 7). The *vraS* and *graR* gene mutations in VISA strains activate arginine catabolism to supply substrates for cell wall biosynthesis (28), while oxidative stress response was triggered to neutralize oxidative damages induced by vancomycin. These metabolic alterations subsequently impose a fitness burden on VISA cells, causing a trade-off between bacterial resistance and virulence.

**Figure 7:**
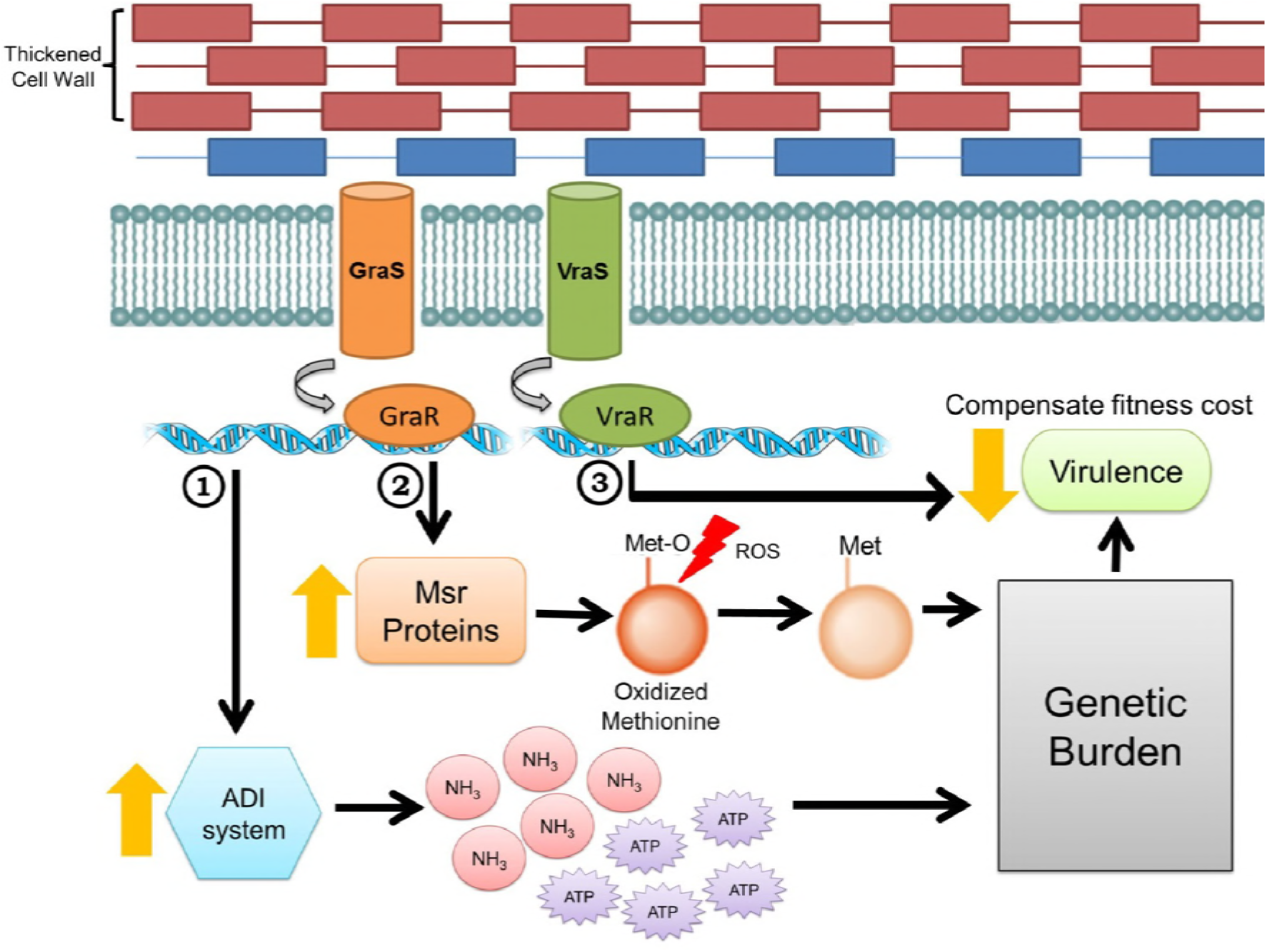
VraSR- and GraSR-mediated regulatory pathways associated with intermediate vancomycin resistance in *Staphylococcus aureus* of the Mu50 lineage: (1) contribution of arginine catabolism (arginine deiminase, ADI) pathway to cell wall thickening, (2) MsrB-associated oxidative stress resistance, and (3) fitness-compensatory response.

## Acknowledgement

The research was funded by the Ministry of Higher Education of Malaysia under the grant codes FRGS/1/2011/SKK/UKM/03/15 and FRGS/1/2014/SKK04/UKM/03/1. We thank Tee Ling Fei and Rosniza Mohamad Hussain for their guidance in handling and maintenance of *C. elegans*. The authors declare no conflict of interest.

## References

1. Beceiro A, Tomás M, Bou G. 2013. Antimicrobial resistance and virulence: a successful or deleterious association in the bacterial world? Clin Microbiol Rev 26:185–230.

2. Pozzi C, Waters EM, Rudkin JK, Schaeffer CR, Lohan AJ, Tong P, Loftus BJ, Pier GB, Fey PD, Massey RC, O’Gara JP. 2012. Methicillin resistance alters the biofilm phenotype and attenuates virulence in *Staphylococcus aureus* device-associated infections. PLoS Pathog 8:e1002626–e1002640.

3. Rudkin JK, Edwards AM, Bowden MG, Brown EL, Pozzi C, Waters EM, Chan WC, Williams P, O’Gara JP, Massey RC. 2012. Methicillin resistance reduces the virulence of healthcare-associated methicillin-resistant *Staphylococcus aureus* by interfering with the *agr* quorum sensing system. J Infect Dis 205:798–806.

4. Peleg AY, Monga D, Pillai S, Mylonakis E, Moellering RC, Jr., Eliopoulos GM. 2009. Reduced susceptibility to vancomycin influences pathogenicity in *Staphylococcus aureus* infection. J Infect Dis 199:532–536.

5. Majcherczyk PA, Barblan J-L, Moreillon P, Entenza JM. 2008. Development of glycopeptide-intermediate resistance by *Staphylococcus aureus* leads to attenuated infectivity in a rat model of endocarditis. Microb Pathog 45:408–414.

6. Kohanski MA, Dwyer DJ, Hayete B, Lawrence CA, Collins JJ. 2007. A common mechanism of cellular death induced by bactericidal antibiotics. Cell 130:797–810.

7. Brynildsen MP, Winkler JA, Spina CS, MacDonald IC, Collins JJ. 2013. Potentiating antibacterial activity by predictably enhancing endogenous microbial ROS production. Nat Biotechnol 31:160–165.

8. Dwyer DJ, Belenky PA, Yang JH, MacDonald IC, Martell JD, Takahashi N, Chan CTY, Lobritz MA, Braff D, Schwarz EG, Ye JD, Pati M, Vercruysse M, Ralifo PS, Allison KR, Khalil AS, Ting AY, Walker GC, Collins JJ. 2014. Antibiotics induce redox-related physiological alterations as part of their lethality. Proc Natl Acad Sci 111:E2100–E2109.

9. Marathe SA, Kumar R, Ajitkumar P, Nagaraja V, Chakravortty D. 2013. Curcumin reduces the antimicrobial activity of ciprofloxacin against *Salmonella typhimurium* and *Salmonella typhi*. J Antimicrob Chemother 68:139–152.

10. Liu Y, Liu X, Qu Y, Wang X, Li L, Zhao X. 2012. Inhibitors of reactive oxygen species accumulation delay and/or reduce the lethality of several antistaphylococcal agents. Antimicrob Agents Chemother 56:6048–6050.

11. Wang X, Zhao X, Malik M, Drlica K. 2010. Contribution of reactive oxygen species to pathways of quinolone-mediated bacterial cell death. J Antimicrob Chemother 65:520–524.

12. Wang X, Zhao X. 2009. Contribution of oxidative damage to antimicrobial lethality. Antimicrob Agents Chemother 53:1395–1402.

13. Tan X-E, Neoh H-m, Looi M-L, Tan TL, Hussin S, Cui L, Hiramatsu K, Jamal R. 2016. Comparative proteomics profiling reveals down-regulation of *Staphylococcus aureus* virulence in achieving intermediate vancomycin resistance. Malays J Microbiol 12:498–505.

14. Romsang A, Atichartpongkul S, Trinachartvanit W, Vattanaviboon P, Mongkolsuk S. 2013. Gene expression and physiological role of *Pseudomonas aeruginosa* methionine sulfoxide reductases during oxidative stress. J Bacteriol 195:3299–3308.

15. Zhao C, Hartke A, La Sorda M, Posteraro B, Laplace J-M, Auffray Y, Sanguinetti M. 2010. Role of methionine sulfoxide reductases A and B of *Enterococcus faecalis* in oxidative stress and virulence. Infect Immun 78:3889–3897.

16. Lee WL, Gold B, Darby C, Brot N, Jiang X, de Carvalho LPS, Wellner D, St John G, Jacobs WR, Jr., Nathan C. 2009. *Mycobacterium tuberculosis* expresses methionine sulphoxide reductases A and B that protect from killing by nitrite and hypochlorite. Mol Microbiol 71:583–593.

17. Atack JM, Kelly DJ. 2008. Contribution of the stereospecific methionine sulphoxide reductases MsrA and MsrB to oxidative and nitrosative stress resistance in the food-borne pathogen *Campylobacter jejuni*. Microbiology 154:2219–2230.

18. Oien DB, Moskovitz J. 2008. Substrates of the methionine sulfoxide reductase system and their physiological relevance, p 94–135. In Schatten GP (ed), Current topics in developmental biology, vol 80. Academic Press, London, UK.

19. Pang YY, Schwartz J, Bloomberg S, Boyd JM, Horswill AR, Nauseef WM. 2014. Methionine sulfoxide reductases protect against oxidative stress in *Staphylococcus aureus* encountering exogenous oxidants and human neutrophils. J Innate Immun 6:353–364.

20. Liu C-I, Liu GY, Song Y, Yin F, Hensler ME, Jeng W-Y, Nizet V, Wang AH-J, Oldfield E. 2008. A cholesterol biosynthesis inhibitor blocks *Staphylococcus aureus* virulence. Science 319:1391–1394.

21. Liu GY, Essex A, Buchanan JT, Datta V, Hoffman HM, Bastian JF, Fierer J, Nizet V. 2005. *Staphylococcus aureus* golden pigment impairs neutrophil killing and promotes virulence through its antioxidant activity. J Exp Med 202:209–215.

22. Singh VK, Vaish M, Johansson TR, Baum KR, Ring RP, Singh S, Shukla SK, Moskovitz J. 2015. Significance of four methionine sulfoxide reductases in *Staphylococcus aureus*. PLoS One 10:e0117594–e0117613.

23. Sifri CD, Begun J, Ausubel FM, Calderwood SB. 2003. *Caenorhabditis elegans* as a model host for *Staphylococcus aureus* pathogenesis. Infect Immun 71:2208–2217.

24. Pukkila-Worley R, Ausubel FM, Mylonakis E. 2011. *Candida albicans* infection of *Caenorhabditis elegans* induces antifungal immune defenses. PLoS Pathog 7:e1002074–e1002086.

25. Irazoqui JE, Troemel ER, Feinbaum RL, Luhachack LG, Cezairliyan BO, Ausubel FM. 2010. Distinct pathogenesis and host responses during infection of *C. elegans* by *P. aeruginosa* and *S. aureus*. PLoS Pathog 6:e1000982–e1001005.

26. Bekeredjian-Ding I, Inamura S, Giese T, Moll H, Endres S, Sing A, Zähringer U, Hartmann G. 2007. *Staphylococcus aureus* protein A triggers T cell-independent B cell proliferation by sensitizing B cells for TLR2 ligands. J Immunol 178:2803–2812.

27. Chávez V, Mohri-Shiomi A, Maadani A, Vega LA, Garsin DA. 2007. Oxidative stress enzymes are required for DAF-16-mediated immunity due to generation of reactive oxygen species by *Caenorhabditis elegans*. Genetics 176:1567–1577.

28. Tan X-E, Neoh H-m, Looi M-L, Chin SF, Cui L, Hiramatsu K, Hussin S, Jamal R. 2017. Activated ADI pathway: the initiator of intermediate vancomycin resistance in *Staphylococcus aureus*. Can J Microbiol 63:260–264.

